# Striatal dopamine synthesis capacity reflects smartphone social activity

**DOI:** 10.1101/2020.06.06.137976

**Authors:** Andrew Westbrook, Arko Ghosh, Ruben van den Bosch, Roshan Cools

## Abstract

Striatal dopamine has been implicated in social behavior across humans, rodents, and non-human primates in artificial laboratory settings with highly-practiced tasks and fixed reward contingencies. Whether striatal dopamine drives naturalistic, spontaneous social behavior remains unclear. Here, we leverage day-to-day logs of unconstrained smartphone behavior and establish a novel link between smartphone social activity and individual differences in striatal dopamine synthesis capacity using [^18^F]-DOPA PET in (N=22) healthy adult humans. We find a strong relationship such that a higher proportion of social app interactions correlates with lower dopamine synthesis capacity in the bi-lateral putamen. Permutation tests and penalized regressions provide evidence that this link between dopamine synthesis capacity and social versus non-social smartphone taps is specific. These observations provide a key empirical grounding for current speculations about dopamine’s role in digital social behavior.

## Introduction

Striatal dopamine has been implicated in social behavior in artificial laboratory settings in humans and animals. The relationship between striatal dopamine function and naturalistic social behavior, however, remains unclear. Smartphones offer a powerful tool for examining spontaneous, real-world social behavior. Their ubiquity has prompted speculation about how smartphone use reflects individual differences in social engagement and the cognitive, affective, and neural systems supporting social interaction (1-3). For example, brain volumetric analyses reveal negative correlations between nucleus accumbens grey matter and both self-reported social media use (4) and the frequency of daily smartphone social app usage (5).

While smartphone behavior has largely been examined via self-report, it is possible to capture richer, more reliable measures by directly monitoring smartphone use (6). In this study, we recorded touchscreen interactions to assess a relationship between social behavior – proxied by the app in use – and striatal dopamine function. Specifically, healthy young adult participants (N = 22; ages 18 to 33; 9 women) completed a ([^18^F]-DOPA) PET scan to quantify their striatal dopamine synthesis capacity. We then examined touchscreen logs over 2—3 weeks of normal, daily use. Following a prior study linking smartphone use measures to motor variability (6), our analysis focused on overall usage (interactions per day) and proportion of social interactions (proportion of all interactions that occurred on social apps – e.g. chat and messenger apps). Given prior work linking striatal dopamine function and motoric vigor (7), we also analyzed peak daily smartphone interaction speed (fastest inter-touch intervals performed every day).

## Results

After aligning participants’ PET scans, we performed a voxel-wise multiple regression to ask where each of these three measures reflected individual differences in dopamine synthesis capacity. The fitted model (Fig. 1A) revealed positive regression weights of striatal dopamine synthesis capacity on our proxy of vigor (interaction speed), negative weights on interactions per day, and negative weights on the proportion of social interactions.

**Fig. 1.**
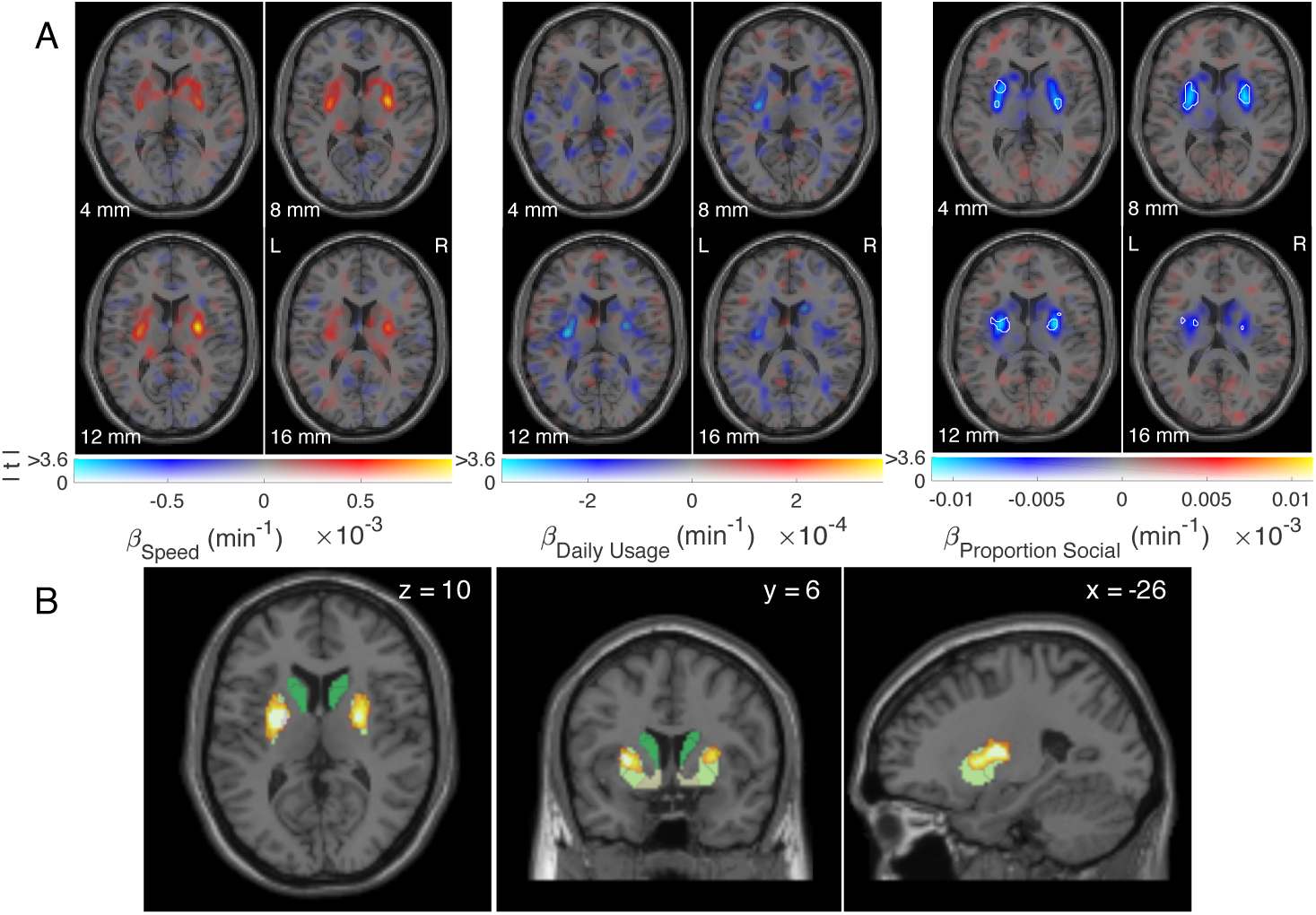
(A) Whole-brain individual difference regression weights of dopamine synthesis capacity predicted by smartphone interaction speed, interactions per day, and proportion of social interactions. White boundaries encompass voxels at (uncorrected) *P* < 0.001, indicating that bi-lateral putamen synthesis capacity reflects proportion of social interactions; display format from (8). (B) Proportion social interaction clusters (*P*_FWE_ < 0.05, small volume correction) resulting from permutation-based, threshold free cluster enhancement (9). Clusters overlap an independently-defined segmentation of the striatum including the dorsal and medial caudate nucleus (dark green), anterior and posterior putamen (yellow-green), and ventral striatum (pale yellow). Note that similar clusters were obtained following thresholded (*P* < 0.001) cluster formation (see Fig. S2).

The negative relationship between the proportion of social interactions and dopamine synthesis capacity was reliable in bi-lateral posterior putamen. Both threshold-free (Fig. 1B) and thresholded cluster forming methods (Fig. S2) confirmed the reliability of the relationship in the putamen in both hemispheres. We restricted our search to a dopamine-rich region encompassing the midbrain and striatum (Fig. S1), and thus retained clusters falling below threshold of *P*_FWE_ < 0.05, small volume corrected.

To probe the relationship with the proportion of social interactions, we next extracted mean dopamine synthesis capacity value from all voxels within the bi-lateral posterior putamen as defined by an independent, functional connectivity-based parcellation of the striatum (10). A robust multiple regression confirmed a negative relationship between individuals’ mean dopamine synthesis capacity and the z-scored proportion of their smartphone interactions devoted to social apps (Fig. 2A; *β* = -1.5×10^−3^ min^-1^, *t*(18) = -4.8, *P* = 1.3×10^−4^) surviving correction for multiple comparisons across the five independently-defined striatal sub-regions (*P*_Bonferroni_ = 6.5×10^−4^). Permutation testing confirmed that the slope of the relationship between dopamine synthesis capacity and social app categorization was extreme compared to 1000 random permutations of social versus non-social app category labels (Fig. 2B). Neither speed (*β* = 7.9×10^−4^ min^-1^, *t*(18) = 2.3, *P* = 0.031), nor overall smartphone usage (*β* = -3.2×10^−4^ min^-1^, *t*(18) = -1.0, *P* = 0.32) relate reliably to dopamine synthesis capacity in the bi-lateral posterior putamen. Collectively, this multiple regression model explains considerable between-subjects variance in posterior putamen dopamine synthesis capacity (*R*^2^ = 0.55).

**Fig. 2.**
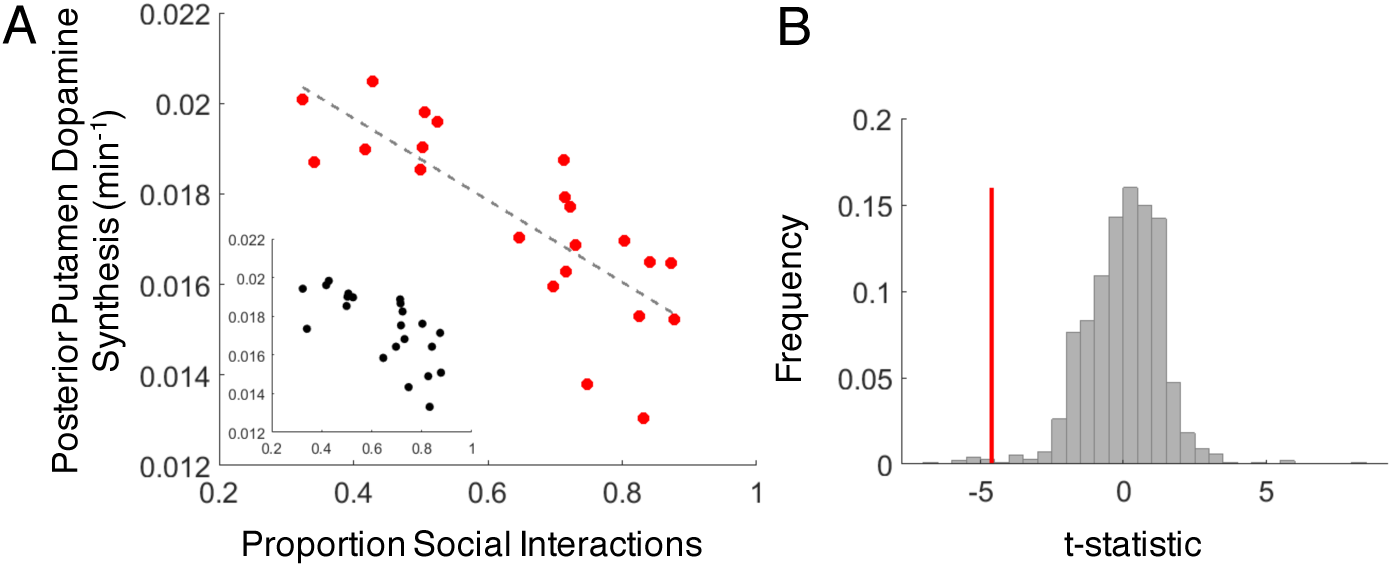
(A) Inverse relationship between individual differences in proportion social interactions and dopamine synthesis capacity (adjusted response plot of the multiple regression including speed and daily usage; inset shows unadjusted data). Dashed line is the estimated linear fit. (B) The observed multiple regression slope (red) is extreme compared to the relationship observed in on 1000 random permutations of app category labels (grey).

A key feature of social activity is that it relies heavily on text messaging, raising the possibility that the observed relationship reflected intensive keypad typing rather than social interactions. However, several non-social apps (e.g., Notes) also rely on typing. Therefore, we used a new category of ‘typing’ apps to control for this effect. We added typing apps – along with age and gender of the participant – as explanatory variables in a lasso regression for parameter selection. Social activity survived the lasso regression (1000 bootstraps; Fig. S3), suggesting that the amount of typing per se cannot explain the link between social activity and dopamine synthesis capacity. Thus, social interactions track between-subjects variance in dopamine synthesis capacity better than typing, age, and gender.

## Discussion

We find a strong and focal relationship between dopamine synthesis capacity in the bi-lateral putamen and smartphone social app use, that is also specific to social app use. This result implies a role for putamen dopamine in smartphone-based social behavior. This result also highlights the promise of passively-tracked, rich smartphone data. It implies, for example, that smartphone social behavior may be a reliable, inexpensive proxy for striatal dopamine synthesis capacity, which can otherwise be assessed only via PET imaging.

Our result informs speculations about digital social behavior and striatal dopamine function. Consider, for example, problematic social media use. Excessive social media use has been associated with reduced ventral striatal grey matter volume (4, 11), suggesting that problematic digital behavior (as in internet addiction) is driven by aberrant reward processing structures. We find a greater proportion of social app usage among those with lower dopamine synthesis capacity. Our results dovetail, albeit in a healthy sample, with the findings that individuals with ADHD have both lower dopamine synthesis capacity (12, 13) and are more prone to social media addiction (14). The fact that one of these studies showed reduced dopamine function specifically in the putamen in ADHD (13) converges with the locus of our result – although we did not a priori predict social app use to relate to putamen dopamine function over other striatal sub-regions. Interestingly, *higher* rather than *lower* dopamine synthesis capacity has been associated with other behavioral addictions (e.g. gambling (15); though see (16)) and behavioral disinhibition (17) indicating that dopamine synthesis capacity may interact with other factors in conferring risk for problematic social media use. One such factor may be D2 receptor density, which is positively correlated with trait extraversion in healthy adults (18).

In sum, our findings highlight the promise of smartphone behavioral data both for elucidating rich, real world social behavior and the neural mechanisms that support them.

## Materials and Methods

We recruited healthy, young adults in the Radboud University community (Nijmegen, Netherlands) who had completed a PET scan as part of a larger pharmaco-imaging study on the influence of catecholamines on cognitive control. A full list of exclusion criteria and study measures collected for the larger study is registered at https://www.trialregister.nl/trial/5959. In total, N = 27 participants responded that they were willing and able to participate. After giving informed consent, a proprietary app (TapCounter, QuantActions; Lausanne, Switzerland) was installed on their smartphone and activated so that their smartphone use data (interactions) could be passively recorded over an interval of 2-3 weeks, and periodically uploaded to a cloud-based server during this interval. 5 participants were subsequently excluded due to data loss from logging and connectivity issues and thus our final sample size was N = 22.

Smartphone data contained the following raw values: the time-stamp of the touchscreen interaction, the label of the app in use at the time of the interaction, and screen on/off times. From these, we computed our three measures of interest: 1) smartphone usage, 2) proportion of social interactions and 3) interaction speed. Smartphone usage was quantified as the square root-transformed total number of interactions, normalized by the number of days of recording. Interaction speed was estimated as follows: the shortest 25^th^ percentile of the inter-event intervals was accumulated in 24 h bins; the inverse of the median of these accumulated values was used.

App labels were used to determine the proportion of social interactions: number of interactions on social apps versus all interactions. Social apps were defined as apps whose main function was to allow users to communicate with each other (friends or strangers). Furthermore, the app had to facilitate these social interactions through direct messaging, public posts, and/or voice chat, contained a personal profile on the app, enabled development of the social network and enabled sharing of personal or external content (e.g. WhatsApp). Apps which did not match these criteria were classified as non-social (e.g. news or weather apps). See (6) for a full description of app classification criteria and additional example apps classified as social versus non-social. For permutation testing of the three-variable multiple regression model, the social versus non-social labels on the app classification table were randomly shuffled. Towards lasso regression (see below), an additional app category of ‘typing’ was used, which included all apps containing a keyboard as a main feature (but excluding searches).

Dopamine synthesis capacity was measured using the radiotracer [^18^F]-fluoro-DOPA (F-DOPA) and a Siemens mCT PET-CT scanner (4 x 4 mm voxels, 5 mm slice thickness). One hour prior to F-DOPA injection, participants received 150 mg carbidopa to reduce decarboxylase activity and 400 mg entacapone to reduce peripheral COMT activity. About 50 minutes after administration, participants were administered a low dose CT scan used for attenuation correction, followed by a bolus injection of 185 MBq (5 mCi) max F-DOPA into the antecubital vein. Over 89 minutes, we collected 4 1-minute frames, 3 2-minute frames, 3 3-minute frames, and 14 5-minute frames. Data were reconstructed with weighted attenuation correction, time-of-flight correction, correction for scatter, and smoothed with a 3 mm full-width-half-max kernel. Presynaptic dopamine synthesis capacity was quantified per voxel as F-DOPA influx rate (Ki; min^-1^) using Gjedde-Patlak linear graphical analysis (19) for the frames of 24—89 minutes, with cerebellar grey matter as the reference region, which was obtained via FreeSurfer segmentation. Ki maps were spatially normalized to MNI space and smoothed using an 8 mm FWHM Gaussian kernel.

Multiple linear regression was performed using the three smartphone explanatory variables: (a) smartphone usage, (b) speed, and (c) proportion of social interactions versus putamen dopamine synthesis capacity via robust regression using the rlm function (MASS package v7.3-51.4) in R. Lasso regression was implemented in MATLAB (MathWorks, Natick, USA) for parameter selection and used the additional variables of: (d) age, (e) gender, and (f) proportion of typing app interactions. Only those parameters which resulted in coefficients different from 0 based on 0.5 and 99.5 percentile range of the bootstrapped (1000) lasso coefficients were considered as meaningful contributors to the regression.

All study measures including derived smartphone metrics, and whole brain dopamine synthesis capacity maps and scripts for reproducing all analyses are available at the Donders Data Repository: https://data.donders.ru.nl/.

## Acknowledgements

We thank the individuals who participated in this study, as well as L. Hofmans and J. I. Määttä who helped coordinate participant recruitment. **Funding:** NWO VICI Grant, 453-14-015 (2015/01379/VI) to R.C.; Grants from Holcim Stiftung and Velux Stiftung (No. 1283) to A.G., and NIH Grant F32MH115600-01A1 to A.W.

## Supplementary materials for this manuscript include the following

Supplementary results

Figures S1 to S3

## Supplementary results

We identified bi-lateral clusters of the putamen surviving FWE *P* < 0.05 small volume correction for multiple comparisons. For our analyses, the small volume was defined by a mask of dopamine rich regions encompassing the midbrain, brainstem, and the basal ganglia (Fig. S1). The mask was defined from the distribution of dopamine synthesis capacity signal across the full sample from the larger parent study of N = 94 participants receiving a [^18^F]-DOPA PET scan. Specifically, following (20), we defined a volume encompassing all voxels where dopamine synthesis capacity was 3 standard deviations above the mean signal across the whole brain. While dopamine synthesis capacity can be measured outside of this region, the signal is much stronger in the volume than elsewhere and hence we reasoned that any relationship with smartphone behavior would arise within this small volume.

**Fig. S1.**
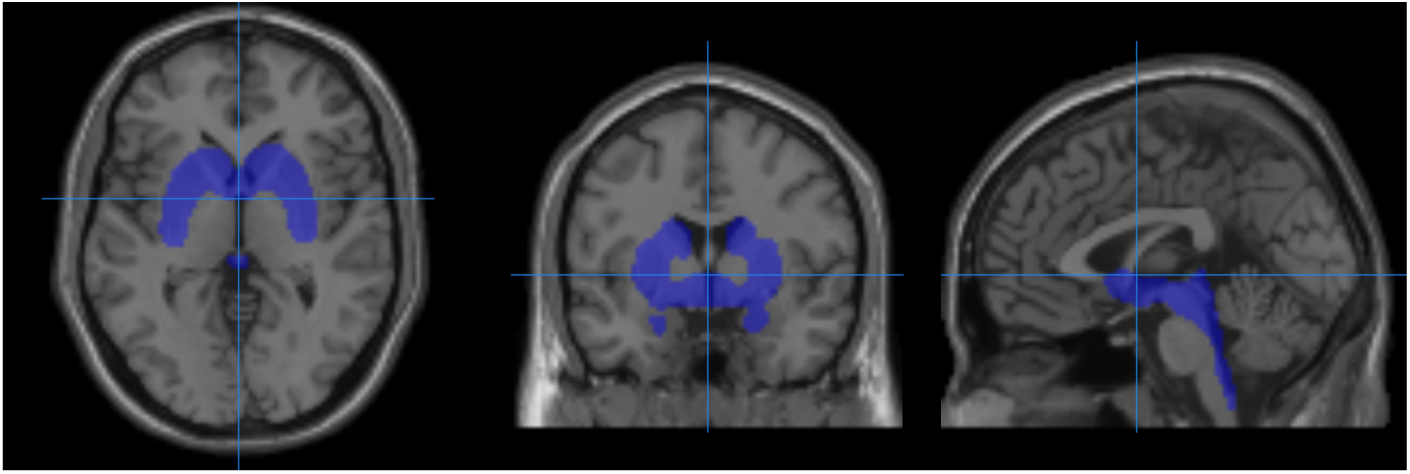
Small volume used for multiple comparisons correction. Volume encompasses all voxels with greater than 3 standard deviations above the mean dopamine synthesis capacity signal across the whole brain. Crosshairs are at MNI [0,0,0].

Complementing our threshold-free cluster enhancement methods in the main text, we also used classic threshold cluster forming methods. We used a seed threshold of *P* < 0.001, with 10 voxels extent, then applied small volume correction and retained clusters surviving FWE at *P*_FWE_ < 0.05. The resulting clusters (Fig. S2) were very similar to those obtained by threshold-free cluster enhancement.

**Fig. S2.**
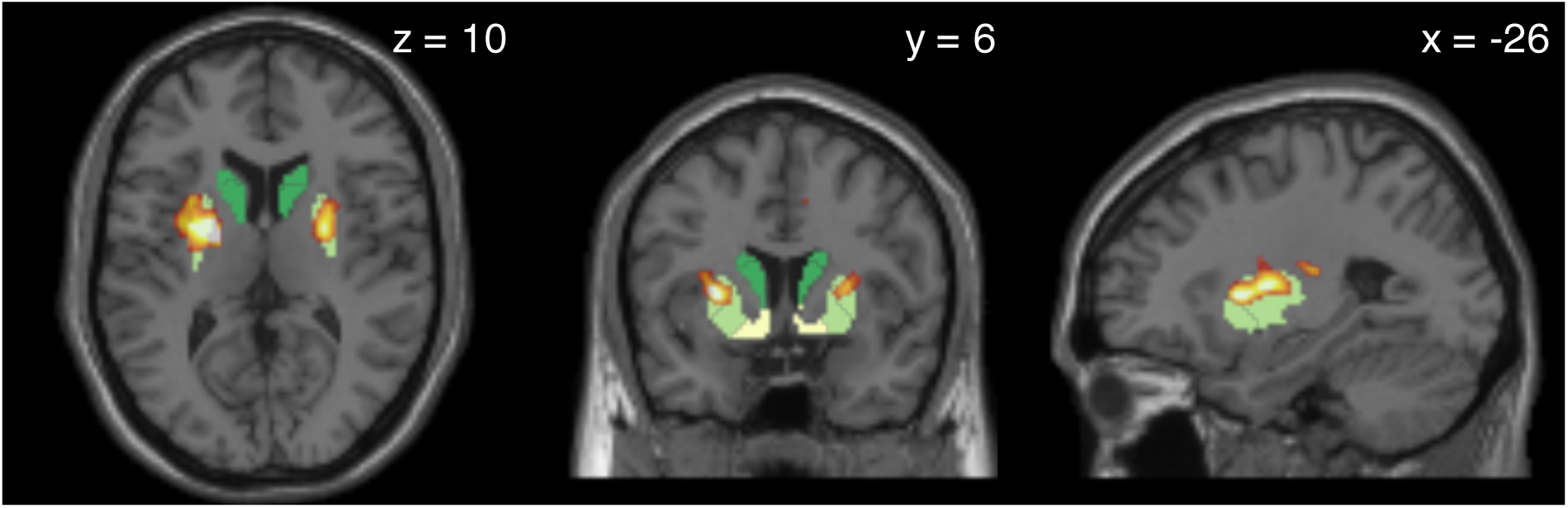
Clusters of voxels showing a reliable relationship between dopamine synthesis capacity and the proportion of social taps following *P* < 0.001 thresholded cluster formation and small volume cluster correction at FWE *P* < 0.05. As in the main text, clusters overlap an independently-defined segmentation of the striatum including the caudate nucleus (dark green), anterior and posterior putamen (yellow-green), and ventral striatum (pale yellow).

While we found a relationship between dopamine synthesis capacity in the posterior putamen and the proportion of touchscreen interactions on social apps, we were also curious whether such a relationship is specific to the posterior putamen and to social app use rather than other potentially explanatory variables. To examine these questions, we extracted mean dopamine synthesis capacity from all voxels in five independently defined regions (10): the anterior (aPut) and posterior putamen (pPut), the medial (mCaud) and dorsal caudate (dCaud), and the ventral striatum (vStr). We then fit lasso regression models, for each of these regions, with the following predictors: total smartphone usage, proportion of social interactions, and interaction speed. We also included the proportion of texting app interactions, participant age and gender. Permutation testing revealed that the proportion of social taps significantly predicts dopamine synthesis capacity in our model of the posterior putamen, but not in other regions (Fig. S3). Moreover, the only reliable predictor in the posterior putamen model was social taps, further demonstrating the specificity of our results.

**Fig. S3.**
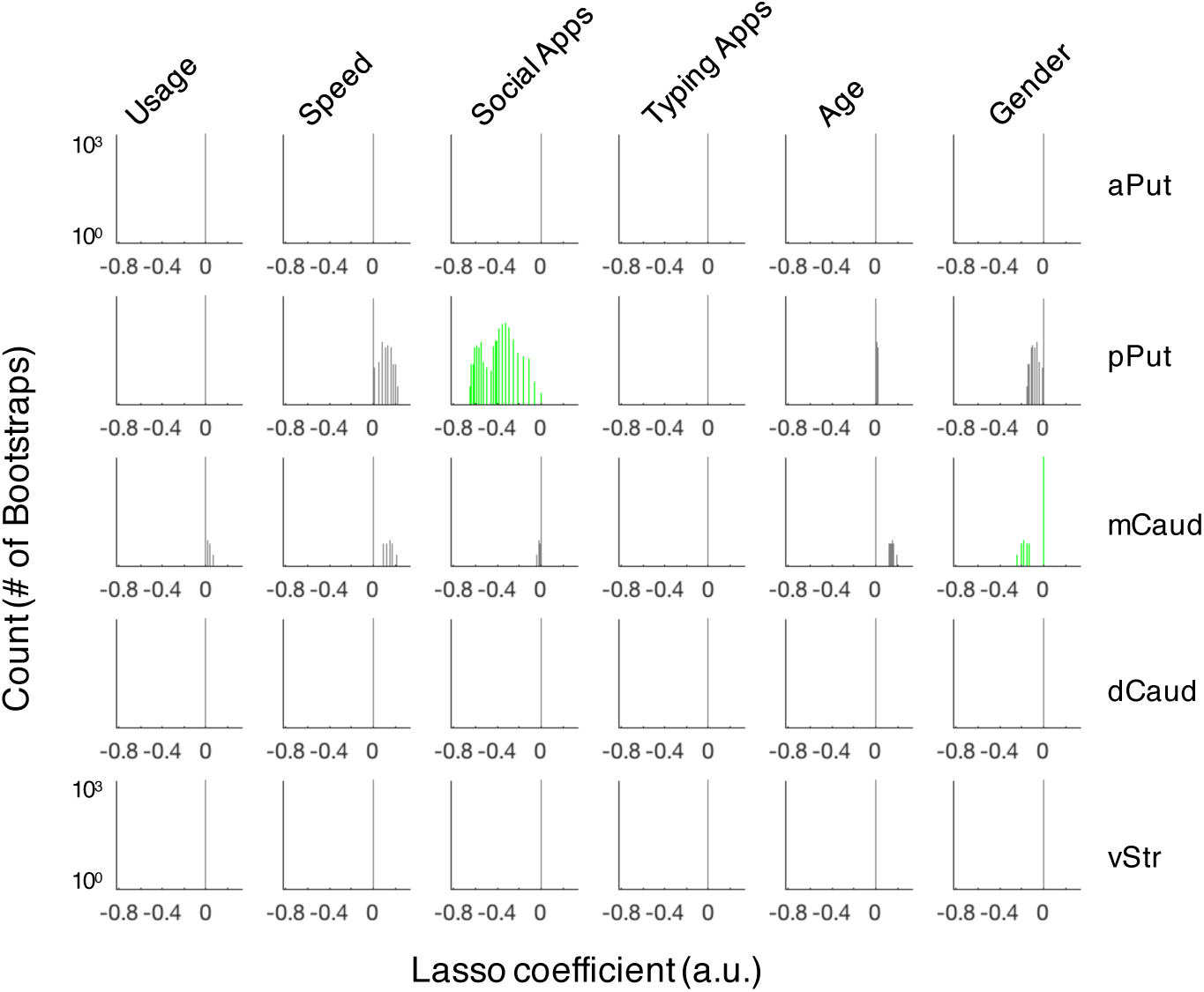
Distributions of lasso regression coefficients testing for relationships between dopamine synthesis capacity in striatal sub-regions and key predictors of interest. Regression coefficients different from 0 based on 0.5 and 99.5 percentile range of the 1000 bootstrapped lasso coefficients are shown in green.

